# hFUT1-based live cell assay to profile α1-2-fucosides enhanced influenza A virus infection

**DOI:** 10.1101/854166

**Authors:** Senlian Hong, Geramie Grande, Chenhua Yu, Digantkumar G. Chapla, Natalie Reigh, Yi Yang, Ken Izumori, Kelley W. Moremen, Jia Xie, Peng Wu

## Abstract

Host cell-surface glycans play critical roles in influenza A virus (IAV) infection ranging from modulation of IAV attachment to membrane fusion and host tropism. Approaches for quick and sensitive profiling of the viral avidity towards a specific type of host-cell glycan can contribute to the understanding of tropism switching among different strains of IAV. In this study, we developed a method based on chemoenzymatic glycan engineering to investigate the possible involvement of α1-2-fucosides in IAV infections. Using a truncated human fucosyltransferase 1 (hFuT1), we were able to create α1-2-linked fucosides *in situ* on the host cell surface to assess their influence on the host cell binding to IAV hemagglutinin and the susceptibility of host cells toward IAV induced killing. We discovered that the newly added α1-2-fucosides on host cells enhanced the infection of several human pandemic IVA subtypes. These findings suggest that glycan epitopes other than sialic aicds should be taken into consideration for assessing the human pandemic risk of this viral pathogen.

---

Belonging to the Orthomyxoviridae family, influenza A viruses (IAV) can circulate widely and cross interspecies barriers.^1^ Their seasonal epidemics at irregular intervals and rapidly evolving nature cause great harm to human health, which makes IAV a moving-target for vaccine and drug development.^2,3^ The IAV life cycle can be divided into five stages: entry into the host cell, entry of the viral ribonucleoprotein (vRNPs) into the nucleus, transcription and replication of the viral genome, the export of the vRNPs from the nucleus; and assembly and budding at the host cell plasma membrane. It is well-known that this cycle is initiated by the binding of hemagglutinin (HA), one of the two key glycoproteins on the outside of the viral envelope, with sialylated glycans on host cells.^4^ Different glycan-binding modes among viral subtypes are common mechanisms that result in the switching tropism.^1,4^ For example, mutations in the glycoreceptor-binding site of HA have been associated with human adaptation of avian IAV, including E190D in the H1 subtype^5^ and Q226L/G228S in the H2 and H3 subtypes^6,7^. Therefore, identifying viral glycan-binding preferences of HA is a priority in order to understand the biology of IVA. Partly due to the lack of tools to mimic the natural presentation of cell-surface diverse glycan epitopes on host cells in a systemic manner, however, limited attention has been drawn to other types of glycans beyond sialylated epitopes.

Besides sialylated glycans, three major types of fucosides are widely expressed in the human respiratory tract, including core α1-6-linked fucosides on N-glycans, and the utmost terminal α1-3-or α1-2-linked fucose.^8–10^ Recent studies have revealed that core-fucosides improve the binding affinity of avian IAV (H1N9) to α2,3-sialylated glycans^11^ and that α1-3-fucosides (e.g., sLe^X^, Neu5Acα2-3Galβ1-4-(Fucα1-3)GlcNAc) bind to human pandemic IVA H1N9 and its HA mutant strains to intensify host cell killing^12^. The relevance of α1-2-fucosides in IAV infection, however, has barely been explored, except a single report that documented that human ‘secretors’ with α1-2-fucosylated epitopes seem to have increased IAV susceptibility compared to ‘non-secretors’ with the fucosyltransferase 2 loss-of-function^13,14^.

To develop a high throughput cell-based assay for profiling the role of α1-2-fucosides in modulating the infection of different IAV strains, we seek to exploit a chemoenzymatic approach for in situ live-cell glycan editing (Fig. 1). In these approaches, the recombinant glycosyltransferases are exploited for *in situ* incorporations of sugar units directly onto the cell-surface glycocalyx by hijacking uncapped glycans to mimic natural multiple presentations of corresponding sugar units in a linkage-specific manner.^15^ Foremost, the full spectrum of live-cell glycans presents natural complexity, in contrast with currently available glycan arrays that none of them are comprehensively contain all the key glycans of live cells, and thus their binding profiles are not predictive of replication.^16,17^ Also, such live-cell based assays has an improved sensitivity for probing the interactions between IAV and a structure-defined glycoepitope, as vividly shown by the hST6Gal1-based live cell assay to differentiate HA-mutants with decreased avidity to the Neu5Acα2-6Gal epitope.^12^

**Figure 1.**
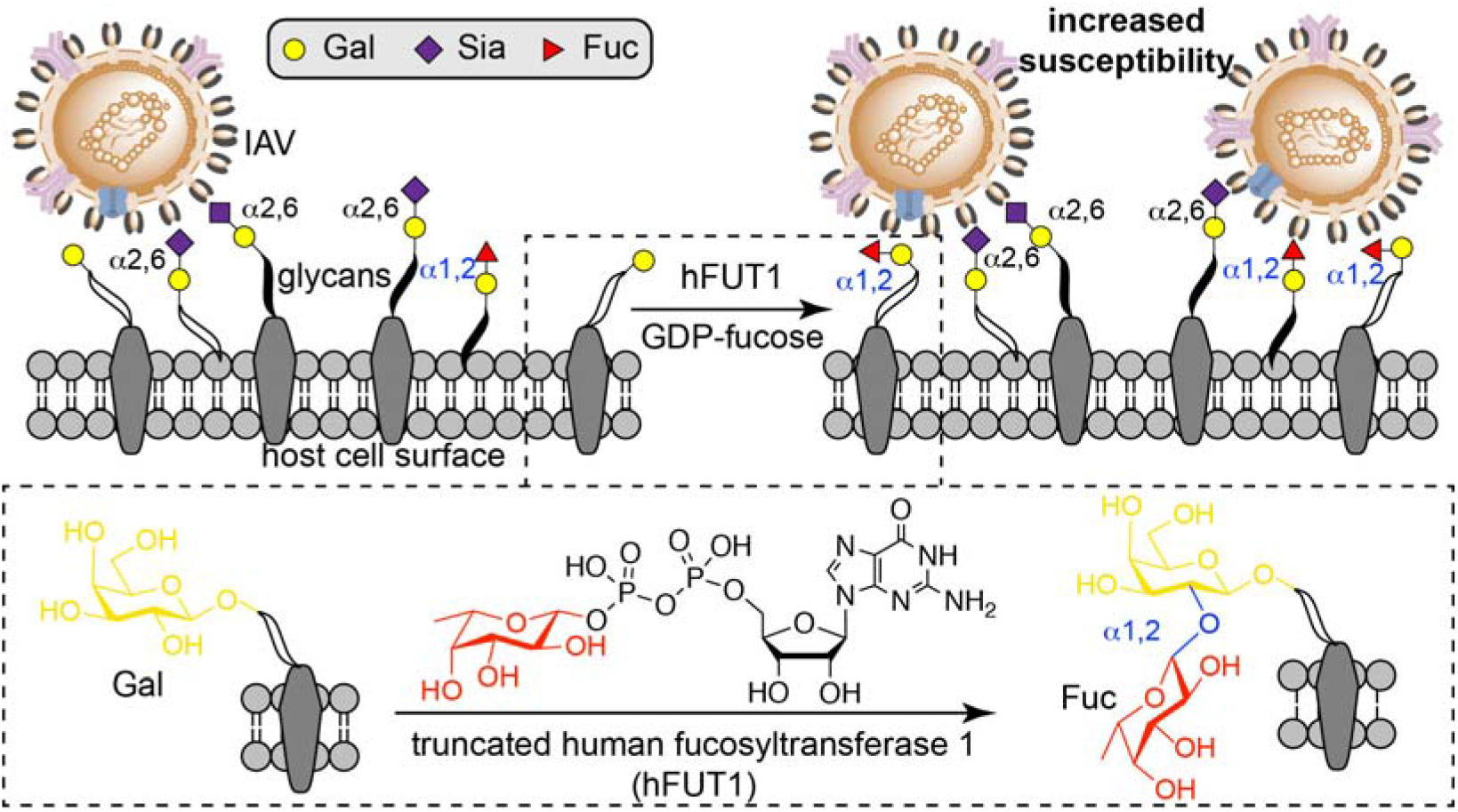
hFUT1-assisted *in situ* creation of α1-2-fucosides on live cells as a novel glycoarray to profile the biological functions of α1-2-fucosides in mediating human pandemic IVA infection. Compared to currently available glycoarrays, cell-based arrays show the full spectrum of live-cell glycans and the additional α1-2-linked fucose, which leads a natural complexity to understand how α1-2-fucosides impact the biology of IAV pathogenesis.

For our specific application, we need an efficient enzyme for *in situ* incorporations of fucose in α1-2-linkages. Recently, the truncated hFUT6^18,19^ has been evaluated as a remarkable tool for *in situ* glycan engineering to create α1-3-fucosides on live cells, among the 13 types of identified human FUTs^10^. Kizuka and coworkers reported that hFUT1, one of the two human galactosidase 2-α-L-fucosyltransferases that are involved in the biosynthesis of α1-2-fucosides, is a tool for *in vitro* synthesis of Fucα1-2Gal analogs with certain substrates promiscuity for alkynes.^20^ Inspired by these studies, we thought to test the human FUT1 for live-cell α1-2-fucosylation.

First, we tested hFUT1-assisted creation of α1-2-fucosides on live Lec2 cells, using GDP-fucose substrate. As a mutant line of Chinese hamster ovary (CHO) cells that have an inactive CMP-sialic acid transporter,^21–23^ Lec2 cells present N-glycans terminated mainly in Gal that could be hijacked by hFUT1 to create α1-2-fucosides. After incubation with hFUT1 and GDP-fucose, the cell-surface α1-2-fucosides were probed with *Ulex europaeus* agglutinin 1 lectin (UEA-FITC) and quantified by flow cytometry. Encouragingly, a robust fluorescence was detected in hFUT1-treated Lec2 cells in a dose-and time-dependent manner (Fig. 2A, 2B and Fig. s1). Compared to a formerly reported *Helicobacter mustelae* α1-2-fucosyltransferase (Hmα1,2FT),^12^ hFUT1 showed a significantly better activity (about 2.5-fold) for incorporating fucose on Lec2 cells under the same conditions, as indicated in the UEA-staining results (Fig. 2B).

**Figure 2.**
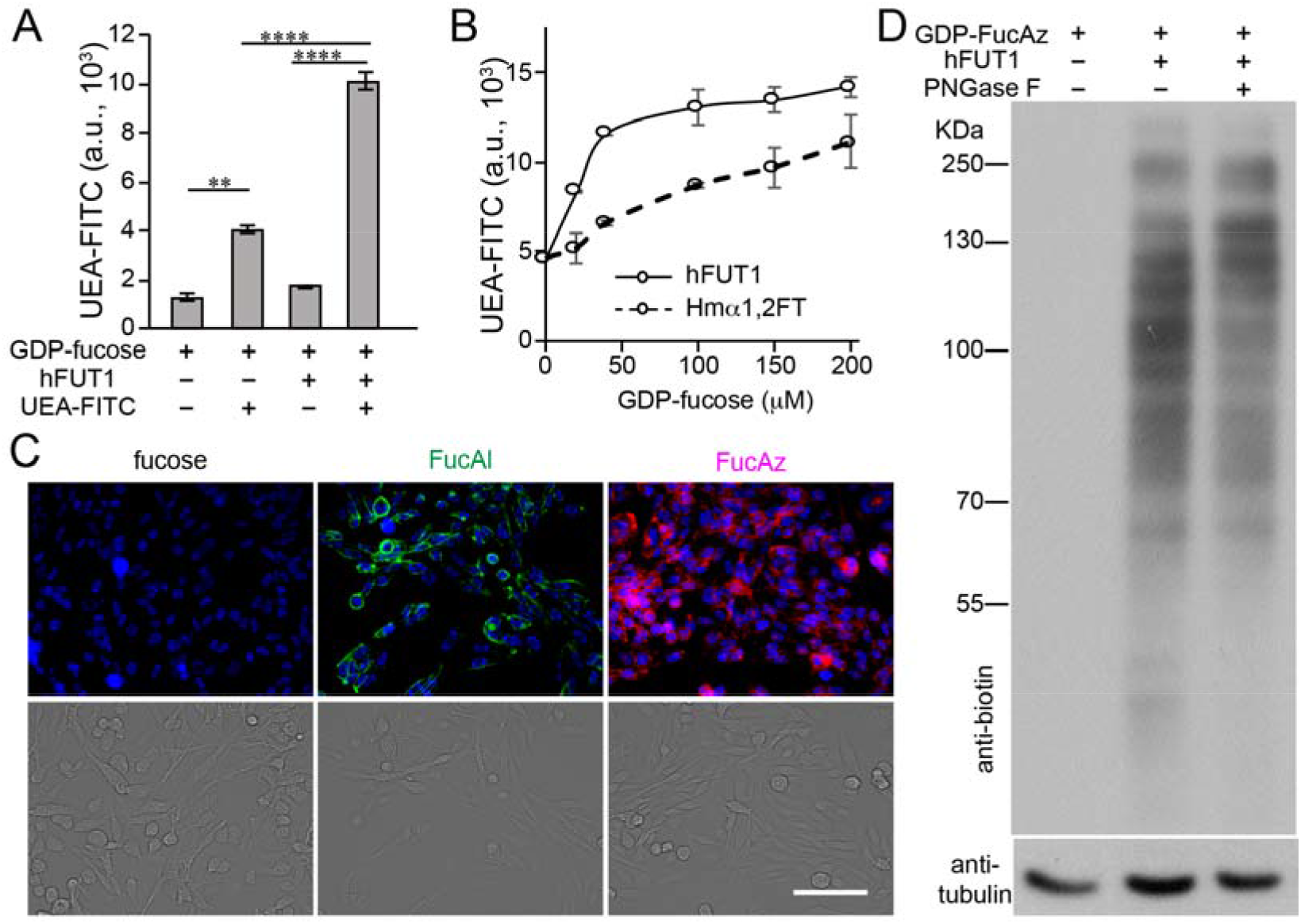
Truncated human fucosyltransferase hFUT1 for live cell-surface glycan engineering. A) Analysis of newly formed α1-2-fucosides on the Lec2 cell-surface by UEA-FITC lectin, the resulting fluorescence was quantified by flow cytometry. B) Comparison of hFUT1 (0.4 mg/mL) and Hmα1,2FT (0.4 mg/mL) assisted creation of α1-2-fucosides on the Lec2 cell-surface under different doses of GDP-fucose. C) Fluorescence microscopy visualization of the incorporated FucAl (Green) or FucAz (Cyan) after the CuAAC-assisted conjugation of biotin, which was further probed with Streptavidin-Alexa Fluor 488. The nucleus was stained with Hoechst 33342. Scale bar, 50 μm. D) hFUT1-assisted incorporation of FucAz onto Lec2 glycoproteins, as detected by anti-biotin western blot (WB). Protein loading was depicted by anti-tubulin WB. In figure A, error bars represent the standard deviation of three biological replicates. ** indicate Student t-test P<0.01; **** indicate Student t-test P<0.001.

Next, we assessed the substrates promiscuity of hFUT1, as good substrate promiscuity would greatly expand the uses of hFUT1. Recent reports have shown the power of glycosyltransferases with excellent substrate promiscuity to incorporate different functionalities to cell-surface glycans, including chemical reporters^15^, affinity tags^24,25^, fluorophores^12^, and biomolecules^25,26^. We first tested the unnatural analogs modified with chemical reporters, including GDP-FucAz and GDP-FucAl.^27^ After incubation with hFUT1 and analogs, the Lec2 cell-surface chemical reporters were further fluorescent-labeled by dyes bearing a complementary group via ligand (BTTPS)-assisted copper catalyzed alkyl-azide [3+2] cycloaddition (CuAAC)^28^. In contrast to the ignorable background signals detected in Lec2 cells treated without hFUT1, strong labeling imaged by fluorescent microscopy suggests that hFUT1 incorporates GDP-FucAl and GDP-FucAz onto Lec2 cell-surface glycans (Fig. 2C). hFUT1-dependent incorporation of azides was further confirmed by dose-dependent incorporation of FucAz (Fig. s2) and the anti-biotin western blot analysis of Lec2 glycoproteins, after the azides were biotinylated via CuAAC for conjugation with Al-PEG4-biotin (Fig. 2D). We also assessed the feasibility of using hFUT1 to incorporate larger substrates,^12^ such as biotinylated-or Cy3 fluorophore conjugated-fucose (Fig. s3A). Unfortunately, only background signals were detected in Lec2 cells treated with GDP-Fuc-biotin or GDP-Fuc-Cy3 compared to the control groups treated without hFUT1 (Fig. s3B), indicating these unnatural donor substrates are not accepted. Compared to Hmα1,2FT that can only use GDP-fucose, hFUT1 enables the incorporation of chemical functionalities—the handles to add additional functionalities.

Taking the efficiency of hFUT1 in creating of α1-2-fucosides, we then used it to develop live-cell based assays to evaluate the role of α1-2-fucosides in the replication cycle of human pediatric IAV. Among the proteins encoded by the eight-segment of the IAV genome, HA is the major surface antigen that mediates initial binding of the viruses to host cell-surface glycoreceptors. To determine if α1-2-fucosides facilitate the binding of HA to host cells, we conducted a binding assay, in which HAs from different viral strains were incubated with Lec2 CHO cells that had been *in situ* modified by hFUT1 to install saturation levels of α1-2-fucosides on the cell surface. Lec2 cells possess a relatively homogeneous *N*-linked glycan profile that does not contain sialylated glycans naturally.^21–23^ Therefore, the role of the newly created α1-2-fucosides can be directly analyzed without the interference of sialic acids.

Using this assay, we tested the HA from A/HongKong/1/ 1968 (HK68) and its laboratory-derived HA-mutant strain HK68-MTA^29^. HK68-MTA (G225M/L226T/ S228A) divaricates in the receptor-binding site (RBS) 220-loop of HA from the wild-type virus (HK68-GLS). Although the wild-type HA-HK68 bound to modified and un-modified Lec2 cells similarly, stronger HA-MTA binding to Lec2 cells modified with α1-2-fucosides was detected (Fig. 3A). Next, we performed two control experiments, in which we converted Lec2 terminal LacNAc into α2-6-sialyl LacNAc and sLe^x^. Consistent with our previous observations, HA-MTA exhibited decreased binding to Lec2 cells with newly installed α2-6-sialic acids^29^, but increased binding to Lec2 cells modified with sialyl Lewis X (sLe^X^)^12^. These observations suggest that sequence flexibility of HK68-HA is capable to commit such viral α1-2-fucoside preferences.

**Figure 3.**
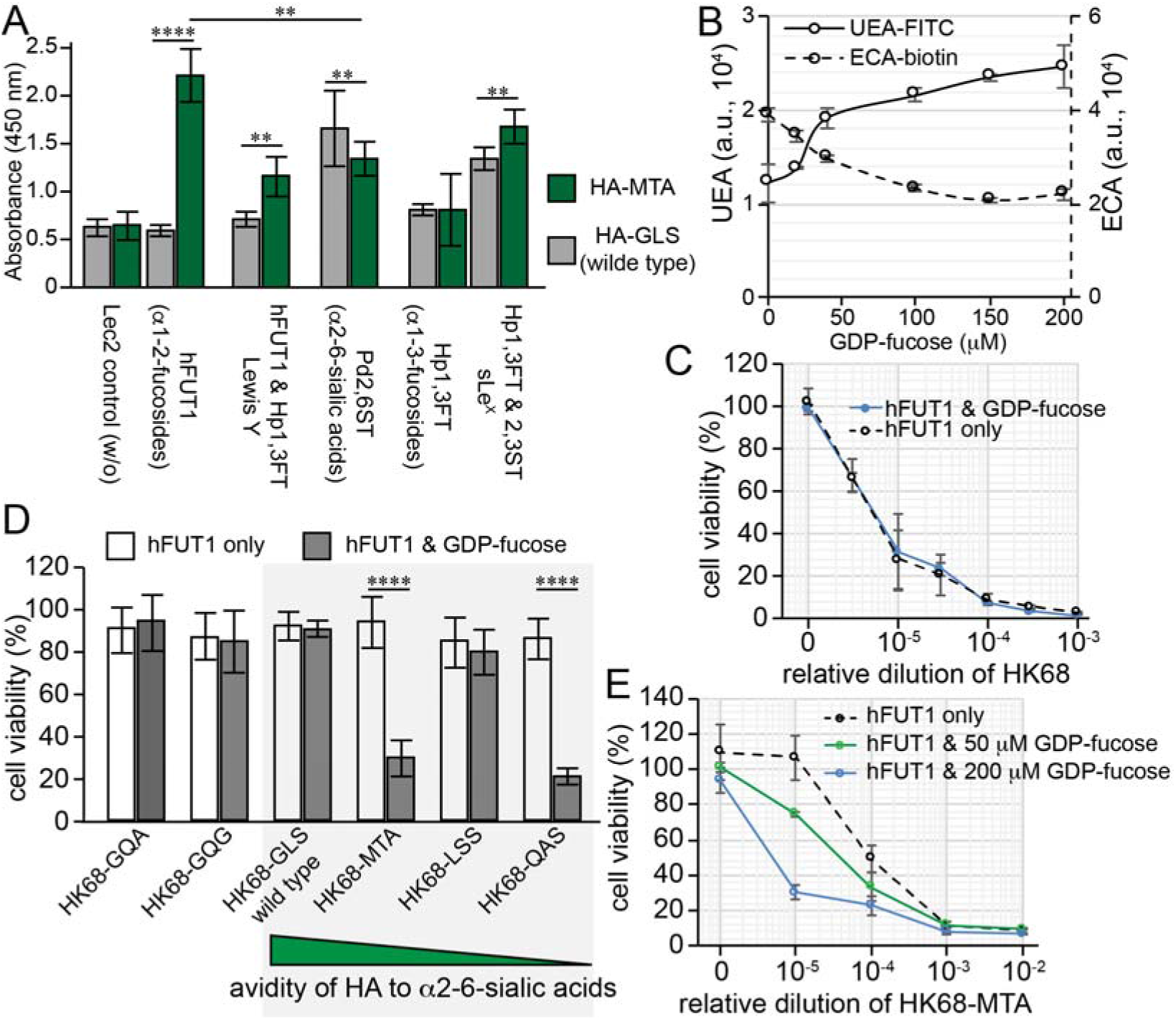
Profiling the structural constraints of IAV-HA for α1-2-fucosides-dependent initial binding and viral infection. The activities of wild-type HK68 and its HA-RBS mutants to infect hFUT1-engineered host cells were directly compared via host cell killing and HA binding assays. A) Relative binding affinity of HA from HK68 wild type (HA-GLS) and HK68-MTA (HA-MTA) for glycan-modified Lec2 cells using the specified recombinant glycosyltransferases. B) Modification of MDCK cell-surface glycocalyx with hFUT1 and GDP-fucose. The α1-2-fucosides on MDCK cell surface was probed using UEA-FITC lectin, and further confirmed via ECA-biotin staining (lectin specific for terminal Gal). C-E) Viability of hFUT1-engineered MDCK cells or control cells upon infection by wild-type HK68 (HK68-GLS) and its HA-RBS mutants, including HK68-QA, HK68-GQG, HK68-MTA, HK68-LSS, and HK68-QAS. In figures A, C-E, the error bars represent the standard deviation of six biological replicates. In figures B, the error bars represent the standard deviation of three biological replicates. ** indicate Student t-test P<0.01; **** indicate Student t-test P<0.001.

Then, we developed an assay to assess if the newly added α1-2-fucosides have any impact on the IAV-mediated host killing. Here, Madin-Darby canine kidney (MDCK) cells, a well-established host cell line for studying IAV, were treated with hFUT1 to create α1-2-fucosylated epitopes. The modified MDCK cells were then infected and analyzed for cell viability after 72 hours upon viral infection. Through this essay, we can correlate the level of host cell-surface α1-2-fucosylation with their susceptibility to the killing induced by different viral strains. The hFUT1-dependent creation of α1-2-fucosides on MDCK cells was first confirmed by the dose-and time-dependent increasing of UEA-FITC staining (Fig. 3B and s4), as well as the decrease of *Erythrina Cristagalli* lectin staining (ECA-biotin, a lectin specific for Gal, N-acetyl galactosamine and lactose).

Similar to HA binding results, infection by the wt HK68 was not affected by the addition of α1-2-fucosides on the MDCK cell surface (Fig. 3C). Nevertheless, a significant increase in host cell killing was observed upon HK68-MTA infection after MDCK cells were modified by α1-2-fucosides (Fig. 3D and 3E). The higher the α1-2-fucoside level, the severer the killing (Fig. 3E). Among the tested HA-RBS mutant strains of HK68, except for HK68-QAS (G225Q/L226A), no other strains including HK68-GQA (L225Q/S228A), HK68-GQG (L226G/ S228G), and HK68-LSS (G225L/L226S) exhibited enhanced killing of α1-2-fucosides-modified MDCK cells (Fig. 3D). These results suggest that the sequence diversifications of HA RBS 220-loop may have a different impact on the binding of IAV to host cells bearing α1-2-fucosides (i.e., Fuc α1-2-Gal or Lewis Y shown in Fig. 3A), which accordingly tunes viral infection and host-cell killing.

Beyond HK68, we also assessed the influence of α1-2-fucosides of the host’s susceptibility to the infection by other subtypes of IAV, including other two additional H3N2 strains A/Perth/16/ 2009 (Perth09) and A/Aichi/2/1968 (Aichi68), and H1N1 strains A/WSN/33 (WSN, H1N1), A/Puerto Rico/8/1934 (PR8) and A/Solomon Islands/3/2006 (SI06). Compared with controls in which MDCK cells were treated with hFUT1 only without the addition of GDP-fucose, α1-2-fucoside-modified MDCK cells were significantly more vulnerable to the IAV-induced killing after exposure to SI06, PR8, Perth09 and Aichi68 strains. By contrast, α1-2-fucoside-modified or unmodified MDCK cells exhibited similar susceptibility to the killing mediated by WSN similar to HK68 strain (Fig. 4A-E, and Fig. 3D). In the Lec2-based binding assay, we also observed a significant increase of the HA (SI06 strain) binding to α1-2-fucoside-modified Lec2 cells (Fig. 4F). So far, about 18 subtypes of HA and 11 subtypes of neuraminidase (NA) have been documented, among which only three subtypes (H1, H2, and H3) have been associated with human pandemics, although five other subtypes (H5, H6, H7, H9, and H10) have sporadically emerged in human. These enhanced infections of aforementioned representative IAV strains, suggests that α1-2-fucosides on human cell-surfaces may enhance the IAV infection, which is further consistent with former observations that ‘secretors’ are more susceptible to influenza A virus^14^.

**Figure 4.**
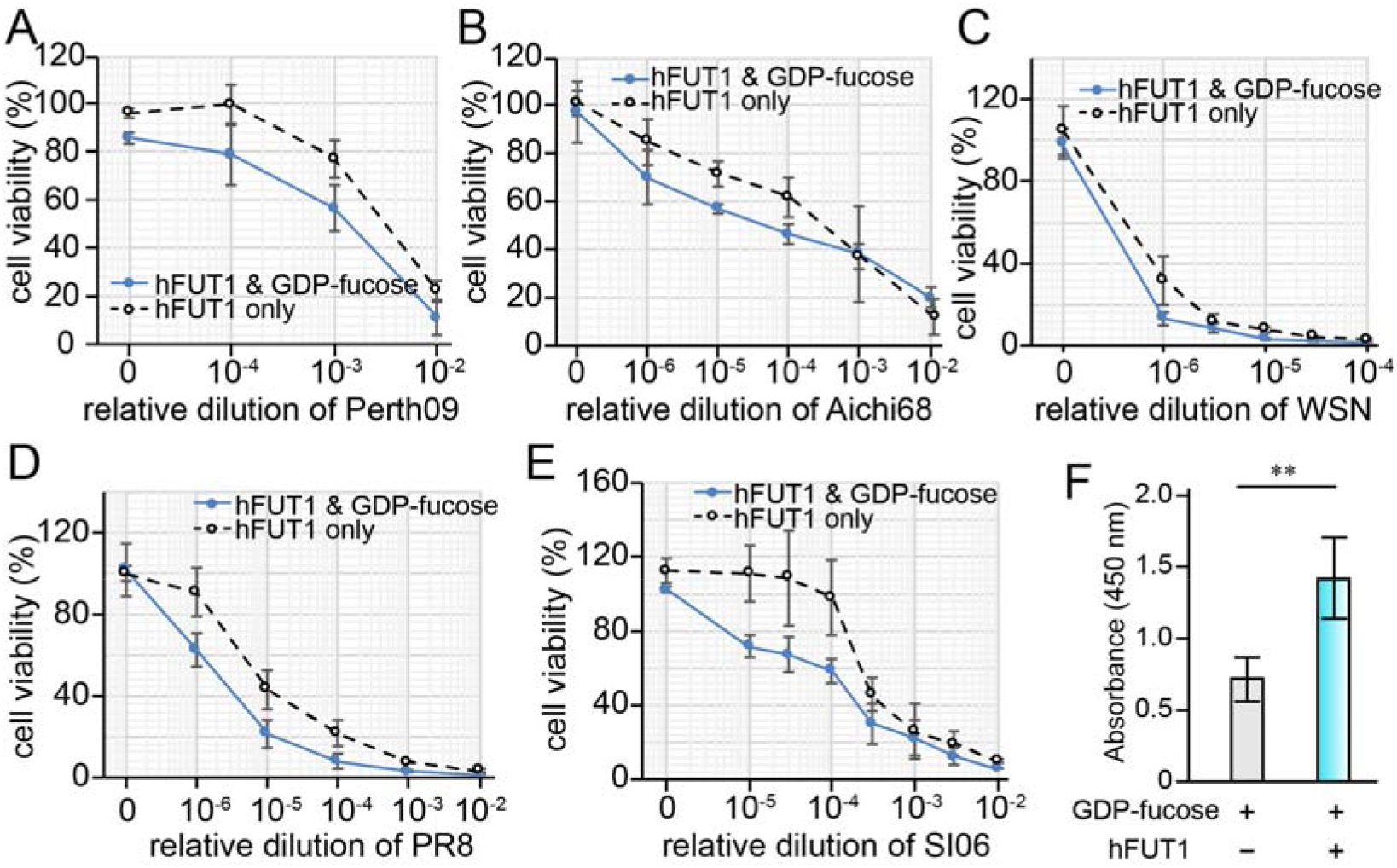
Profiling human pandemic IAV infection with glycocalyx-modified live cells. A-E) cell viability of hFUT1-modified host MDCK or control cells (treated with hFUT1 only) upon infection by Perth09 (A), Aichi68 (B), WSN (C), PR8 (D), SI06 (E) influenza viruses. F) The relative binding affinity of HA from SI06 for α1-2-fucoside-modified Lec2 cells using hFUT1. The error bars represent the standard deviation of six biological replicates. ** indicate Student t-test P<0.01.

Overall, using hFUT1-mediated *in situ* creation of α1-2-linked fucose on live cells, we revealed that the addition of α1-2-fucosylation enhanced the infection of several human pandemic IVA subtypes. The newly created glyco-epitopes (i.e., α1-2-fucosides) may influence many factors in the IAV replication cycle (e.g., initial binding, internalization and viral particle releasing, *etc),* which in turn modulate the IAV-induced host cell killing. Further studies using additional approaches (e.g. virus tracing, neuraminidase-binding and structural analysis, *etc)* are necessary for elucidate the detailed molecular mechanism. Nevertheless, our current study highlights the involvement of glycan epitopes other than sialic acid in influenza virus pathogenesis, which should be taken into consideration for assessing the human pandemic risk of this major pathogen.

